# minute: A MINUTE-ChIP data analysis workflow

**DOI:** 10.1101/2022.03.14.484318

**Authors:** Carmen Navarro, Marcel Martin, Simon Elsässer

**Author notes:** authors contributed equally (Carmen Navarro, Marcel Martin). Correspondence: Simon J Elsässer.

## Abstract

Quantitative ChIP-seq methods are essential for accurately characterizing and comparing genome-wide DNA-protein interactions across samples. Procedures that enable such quantitative comparisons involve addition of spike-in chromatin or recombinant nucleosome material, or a multiplexed process using barcoding of chromatin fragments. ChIP-seq analysis workflows typically require a number of computational steps involving multiple tools in order to reach interpretable results, and quantitative analyses require additional steps that ensure scaling of the processed output according to the quantitative measurements. Crucially, the different quantitative approaches have unique analysis requirements reflecting the disparate experimental workflows, hence no universal analysis pipeline exists for quantitative ChIP-seq. Here, we developed **minute,** a user-friendly computational workflow to easily process multiplexed ChIP data that handles the specific needs of quantitative ChIP. **minute** enables transformation of raw, multiplexed FASTQ files into a set of normalized, scaled bigWig files that can serve as a basis for a quantitative, comparative downstream analysis. **minute** is implemented in Python and Snakemake and paired with a Conda environment, to facilitate usability and reproducibility in different platforms.

Source code of **minute** is available on GitHub: https://github.com/NBISweden/minute

## Background

Data analysis workflow engines are the current standard to ensure reproducible, transparent, quality-controlled research. Reproducible research is an issue that needs to be addressed both from experimentalists (Baker, 2016) and computational biologists (Grüning et al., 2018). Data analyses need to be reproducible and transparent in order to ensure sound scientific research. To this end, many computational frameworks to develop data analysis workflows are available, such as Snakemake (Mölder et al., 2021), Nextflow (Di Tommaso et al., 2017), Galaxy (Giardine et al., 2005), Toil (Vivian et al., 2017). Efforts have been made to provide useful guidelines and engage the life sciences community in this out-of-the-bench side of reproducible science (Shade and Teal, 2015); (Brito et al., 2020) (Jackson et al., 2021). The aforementioned frameworks provide a more expressive, intuitive language to design bioinformatics analyses based on scientific decisions rather than package versions and software environments. Additionally, they are designed to provide better support for desirable features in scientific data analysis, such as reproducibility, re-entrancy, scalability or portability (Perkel, 2019).

Chromatin immunoprecipitation followed by sequencing (ChIP-seq) is an extensively used technology that allows to study location and sequence affinity of DNA-protein interactions (Furey, 2012). There exist many experimental protocols (Furey, 2012) and computational workflows (Nakato and Sakata, 2021) to process ChIP-seq raw data. Standard ChIP-seq experiments have a limited capacity to gauge quantitative differences across samples (Chen et al., 2015; van Galen et al., 2016; Orlando et al., 2014). This restricts the biological conclusions that can be confidently drawn from traditional ChIP-seq experiments to those based on presence or absence of interactions of the target protein, excluding the nuances of scaled, quantitative direct comparison between samples (Chen et al., 2015). Quantitative ChIP-seq methods have been proven to be a valuable approach to accurately measure histone level differences that are overlooked by standard ChIP-seq (van Galen et al., 2016; Kumar and Elsässer, 2019; Kumar et al., 2021; van Mierlo et al., 2019; Orlando et al., 2014). Mint-ChIP (van Galen et al., 2016) and MINUTE-ChIP (Kumar and Elsässer, 2019) are quantitative, multiplexed ChIP-seq methods in which DNA adaptors containing a sample-specific barcode are ligated onto fragmented chromatin before samples are pooled and processed together through the ChIP procedure.

Raw read files produced by MINUTE-ChIP, as much as other traditional ChIP-seq approaches, require a series of quality control, processing and transformation steps that involve multiple computational tools, such as Cutadapt (Martin, 2011), deepTools (Ramírez et al., 2014), samtools (Li et al., 2009). In order to aid automated, reproducible analyses of sequencing data produced by MINUTE-ChIP, we developed **minute** (Multiplexed, Input-Normalized, UMI, T7 ChIP), an open-source bioinformatics pipeline that employs Snakemake (Mölder et al., 2021), a state-of-art workflow engine with a long trajectory and a wide user base. The pipeline is configured with intuitive sample definition files and processes raw or demultiplexed FASTQ files. Resulting output bigWig files are scaled according to input normalization and therefore directly comparable, enabling quantitative downstream genomics analysis using a wealth of existing bigwig suites, R or Python.

## Methods

### Implementation

**minute** pipeline is implemented in Python and Snakemake (Mölder et al., 2021), a workflow management system based on Python programming language. Snakemake workflows can be run on a variety of computing environments, such as local computers, clusters and cloud computing systems. Source code and documentation are available on GitHub: https://github.com/NBISweden/minute. Continuous Integration via GitHub Actions on a test dataset is also implemented to ensure high code reliability. In order to guarantee long-term reproducibility, a Conda (https://conda.io) environment is specified with every tool required to run the **minute** workflow.

### Workflow description

**minute** pipeline’s main function is to transform a set of multiplexed FASTQ files into properly scaled, directly comparable per-sample bigWig files that can be visualized on a genome browser tool such as the Integrative Genomics Viewer (Robinson et al., 2011), and serve as basis for downstream analysis. Additionally, relevant quality control steps are performed and reports are generated throughout the workflow. **minute’s** main steps (see Figure 1) are briefly described below.

**Figure 1.**
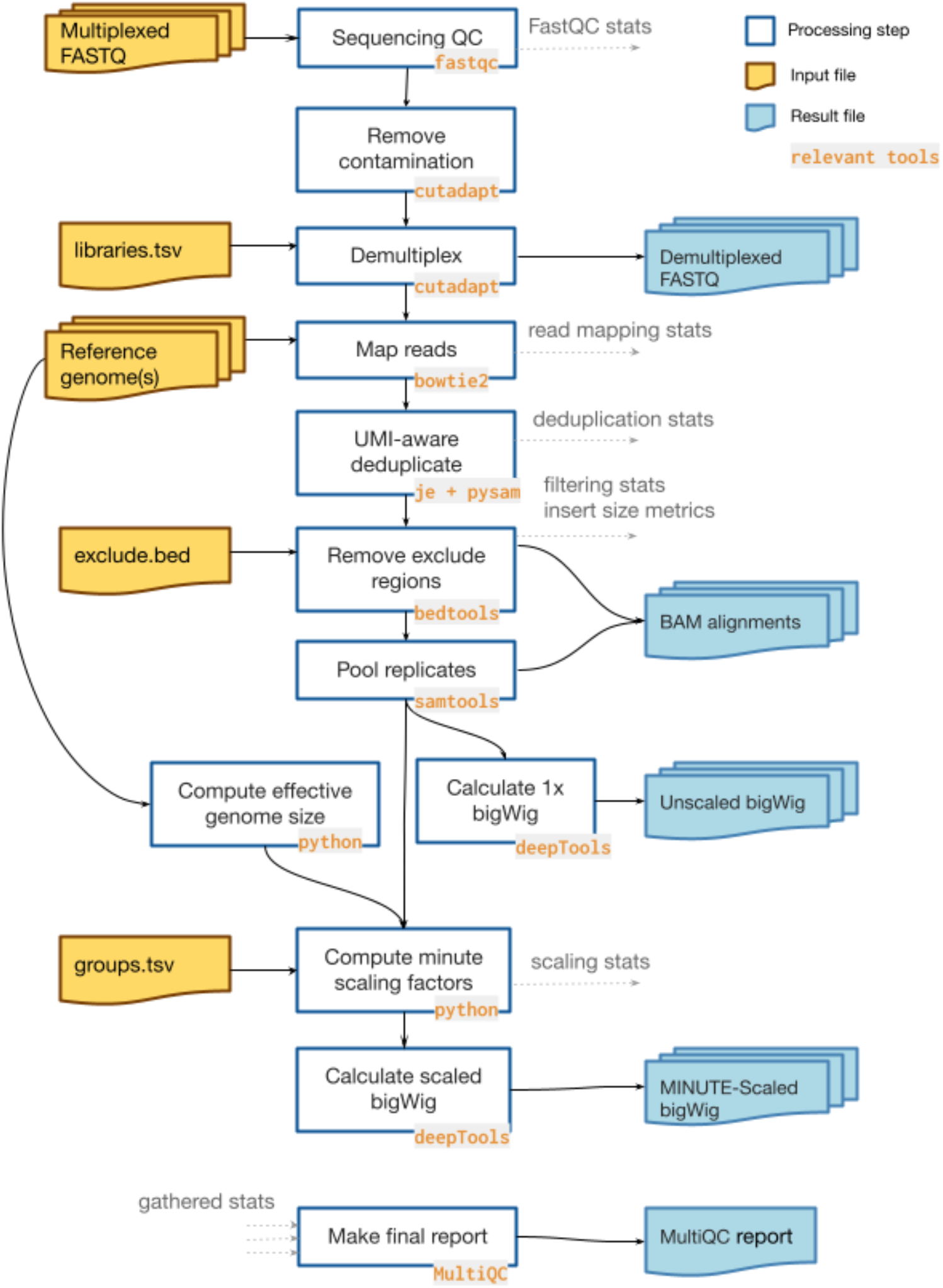
minute pipeline workflow. Yellow files represent necessary inputs. Blue files represent final outputs. Each main processing step is annotated with its corresponding relevant tool. Quality control metrics extracted are shown as annotated grey arrows. Finally, a MultiQC report is generated gathering all QC stats.

#### Input

The main input files needed for **minute** execution are paired-end multiplexed FASTQ files. Additionally, two tables are required: libraries.tsv (see Table 1) specifies the barcode used in each of the multiplexed libraries, and groups.tsv (see Table 2) specifies the scaling configuration and the reference to which each sample should be mapped. An optional BED file containing regions to be filtered out can also be provided. The reference genomes to be used must also be provided as FASTA files with corresponding Bowtie 2 (Langmead and Salzberg, 2012) indexes.

**Table 1.** Example libraries.tsv table content. Column names are included for clarity.

| library | replicate | barcode | fastq_basename |
| --- | --- | --- | --- |
| IN_SL_CTR | 1 | CATGCTTA | testdata-Inp |
| IN_SL_CTR | 2 | GCACATCT | testdata-Inp |
| IN_SL_CTR | 3 | AGTTGCTT | testdata-Inp |
| IN_2i_CTR | 1 | GGTCCAGA | testdata-Inp |
| IN_2i_CTR | 2 | GTATAACA | testdata-Inp |
| IN_2i_CTR | 3 | AGGGTACT | testdata-Inp |
| H3K4m3_SL_CTR | 1 | CATGCTTA | testdata-H3K4m3 |
| H3K4m3_SL_CTR | 2 | GCACATCT | testdata-H3K4m3 |
| H3K4m3_SL_CTR | 3 | AGTTGCTT | testdata-H3K4m3 |
| H3K4m3_2i_CTR | 1 | GGTCCAGA | testdata-H3K4m3 |
| H3K4m3_2i_CTR | 2 | GTATAACA | testdata-H3K4m3 |
| H3K4m3_2i_CTR | 3 | AGGGTACT | testdata-H3K4m3 |

**Table 2.** Example groups.tsv table. For each group, the first library is considered the reference, and rest of libraries are scaled accordingly.

| library | replicate | input | scaling group | reference |
| --- | --- | --- | --- | --- |
| H3K4m3_SL_CTR | pooled | IN_SL_CTR | H3K4m3 | mm9 |
| H3K4m3_SL_CTR | 1 | IN_SL_CTR | H3K4m3 | mm9 |
| H3K4m3_SL_CTR | 2 | IN_SL_CTR | H3K4m3 | mm9 |
| H3K4m3_2i_CTR | pooled | IN_2i_CTR | H3K4m3 | mm9 |
| H3K4m3_2i_CTR | 1 | IN_2i_CTR | H3K4m3 | mm9 |
| H3K4m3_2i_CTR | 2 | IN_2i_CTR | H3K4m3 | mm9 |

#### Quality Control

QC metrics are generated at several steps of the pipeline and their outputs together with each step result are summarized into a final report using MultiQC (Ewels et al.,2016). FASTQ files are processed using FastQC (https://www.bioinformatics.babraham.ac.uk/projects/fastqc/). Insert size metrics are gathered using Picard (https://github.com/broadinstitute/picard), and mapping statistics are generated with SAMtools (Li et al., 2009).

#### Contamination removal

Known contaminant sequences (e.g. MINUTE adaptor concatemers or poly-G second reads from failure of cluster reversal in the Illumina flow cell) are removed explicitly using Cutadapt (Martin, 2011) with an error tolerance of 15%.

#### Demultiplexing

FASTQ files are demultiplexed with Cutadapt (Martin, 2011) using a sample-specific barcode of 8 nucleotides at the 5’ end of read 1 (first read in pair). By default, one mismatch is allowed for assigning each read to its corresponding sample since barcode pools should usually keep a minimum Hamming distance of 3. The allowed number of mismatches can be configured to better fine-tune performance to different barcode pool distributions.

#### Read mapping

Reads are mapped to the specified reference genome using Bowtie 2 (Langmead and Salzberg, 2012) with parameters --reorder (for reproducibility) and --fast. When pooled replicates are specified in the groups table, the generated BAM files are merged into a single one using SAMtools (Li et al., 2009).

#### Deduplication

Duplicate reads are found with je markdupes (https://github.com/gbcs-embl/Je/), which marks reads as duplicate if both their mapping locations and UMIs match. The program is run on a version of the BAM file that only contains the read 1 (first read in pair) sequences since T7-based amplification does not guarantee amplification of the full-length template fragment. Duplicate read pairs are then removed from the original BAM files by a custom script that uses pysam.

#### Remove excluded regions

If configured, reads overlapping regions specified in a custom BED file are removed from the analysis using BEDTools (Quinlan and Hall, 2010). This usually includes loci that have been previously annotated as artifact-prone (Amemiya et al.,2019) and-or repetitive regions depending on the nature of the experiment.

#### Input-normalized scaling factors

Scaling factors are calculated at this step to generate a set of genomic profiles that are quantitatively comparable. The rationale for calculating scaling factors is as follows: in an theoretical ideal scenario, all sample barcodes in the input pool are represented exactly equiproportional. Following from the biochemical binding equilibrium, a library prepared from the ChIPed DNA will then contain a barcode distribution that reflects proportionally the relative abundance of the immunoprecipitated epitope in each of the pooled samples. In practice, barcoding efficiency varies between samples due to experimental imperfection (small variations in cell number, volumes, adaptor oligo amount and reaction conditions). Hence, the input barcode distribution must be separately assessed from an Input library, and the read counts for each barcode in the ChIP library are normalized to their respective counts in the input library, yielding input-normalized ratios (INRs). In an equilibrium binding, INRs will reflect the relative abundance of the immunoprecipitated epitopes *independent* of the input variability, as also validated experimentally (Kumar and Elsässer, 2019).

The scaling factor is calculated as follows:

Let Reads_ChIP,barcode_ be the number of MINUTE-ChIP mapped, deduplicated reads in a given condition (i.e. barcode) for a given ChIP, and Reads_input,barcode_ the corresponding number of MINUTE-ChIP input reads for the same condition. We define the **Input Normalized Ratio** as:

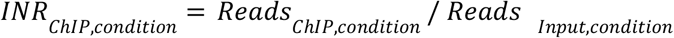
To convert these raw INRs to a intuitively meaningful value, one condition (represented by a single barcode or replicate barcodes) is chosen as the reference condition. The reference condition receives a global mean of 1 (as commonly used in Reads per Genome Coverage, also termed “1x Genome Coverage” normalization methods). The remaining conditions in the same ChIP pool are scaled *relative* to this reference condition. We define the **minute-Scaled Ratio** for such ChIP, reference and condition as:

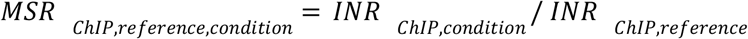
Multiplying this ratio by a 1x genome coverage factor we can calculate the corresponding **minute-Scaled Read Counts**:

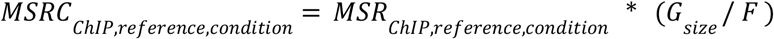
Note that MSRC_ChIP,reference,condition_ = *G_size_/F* when reference = condition, hence MSRC_ChIP,reference,condition_ becomes a constant that represents 1x genome coverage for the corresponding reference genome and fragment length (RPGC).
A factor that scales the BAM file to the final quantitative track is calculated accordingly to generate scaled bigWig files in the next step. The scaling factor K_ChIP,referencei,condition_ is defined therefore as:

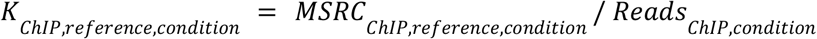
K value is used as --scaleFactor parameter for deepTools when generating each bigWig file in the next step.

#### BigWig generation

minute-scaled bigWig files are produced using deepTools (Ramírez et al., 2014) using --scaleFactor K, calculated as described above. deepTools requires the effective genome size as another parameter and this is calculated as the number of non-N nucleotides in the reference genome provided as input. Unscaled bigWig files disregarding the quantitative scaling information are generated as well, each normalized individually to 1x GenomeCoverage.

#### Execution report

Finally, the **minute** pipeline runs MultiQC (Ewels et al., 2016) to collect statistics generated at each execution step and assembles these into a summary report. The report is customized to include a scaling table and graph.

### Running minute

#### Installation

To install minute, we recommend using Conda and Bioconda:

1. **Install Conda** and Bioconda by following the instructions at httD://bioconda.github.io/user/install.html
2. **Create a Conda environment containing minute**:

~~~
conda create -n minute minute
~~~
3. **Activate the “minute” environment**:

~~~
conda activate minute
~~~
4. **Run minute**:

~~~
minute --help
~~~

### Setup and run

#### Input files

minute pipeline’s main input data are paired-end FASTQ files containing multiplexed samples. Each read 1 starts with a Unique Molecular Identifier (UMI) of user-defined length and is followed by a 8-bp barcode identifying the sample. Alternatively, the files can be already demultiplexed, which ensures that minute can be re-run on already published datasets available from the Gene Expression Omnibus (GEO) database. The demultiplexing information is extracted from the experimental design files described below.

#### Experimental design

Two metadata files are needed:

**`libraries, tsv`**. This is a tab-separated file that contains information about the barcodes and replicates (see Table 1). This file has 4 columns: Sample name, replicate number, barcode identifying the library, and FASTQ file base. If the file is already demultiplexed, a dot (“.”) should be entered instead of the barcode. The FASTQ base name refers to files within the fastq/ folder. The suffixes `_R1.fastq.gz` and `_R2.fastq.gz` will be added automatically.
**`groups, tsv`**. This is a tab-separated file with information about each library’s matching input and scaling groups, including which condition serves as reference per ChIP and which reference genome to use (see Table 2). It is possible to specify different scaling groups, each one with its own reference library. For each scaling group, the first library will be taken as reference. Note that a given library cannot be specified in multiple scaling groups at the same time. This file has 5 columns: sample name (which needs to match one of the sample names in `libraries.tsv`), replicate number (this can also contain the word “pooled” to indicate that all replicates of that sample are to be pooled), corresponding input (matching a library in the `libraries.tsv` file), scaling group name and genome reference. The reference genome must be one of those specified in the `minute.yaml` file.

#### Run configuration

`minute.yaml` specifies the reference genomes used, excluded loci and other configuration values, such as UMI length (Figure 3).

**Figure 2:**
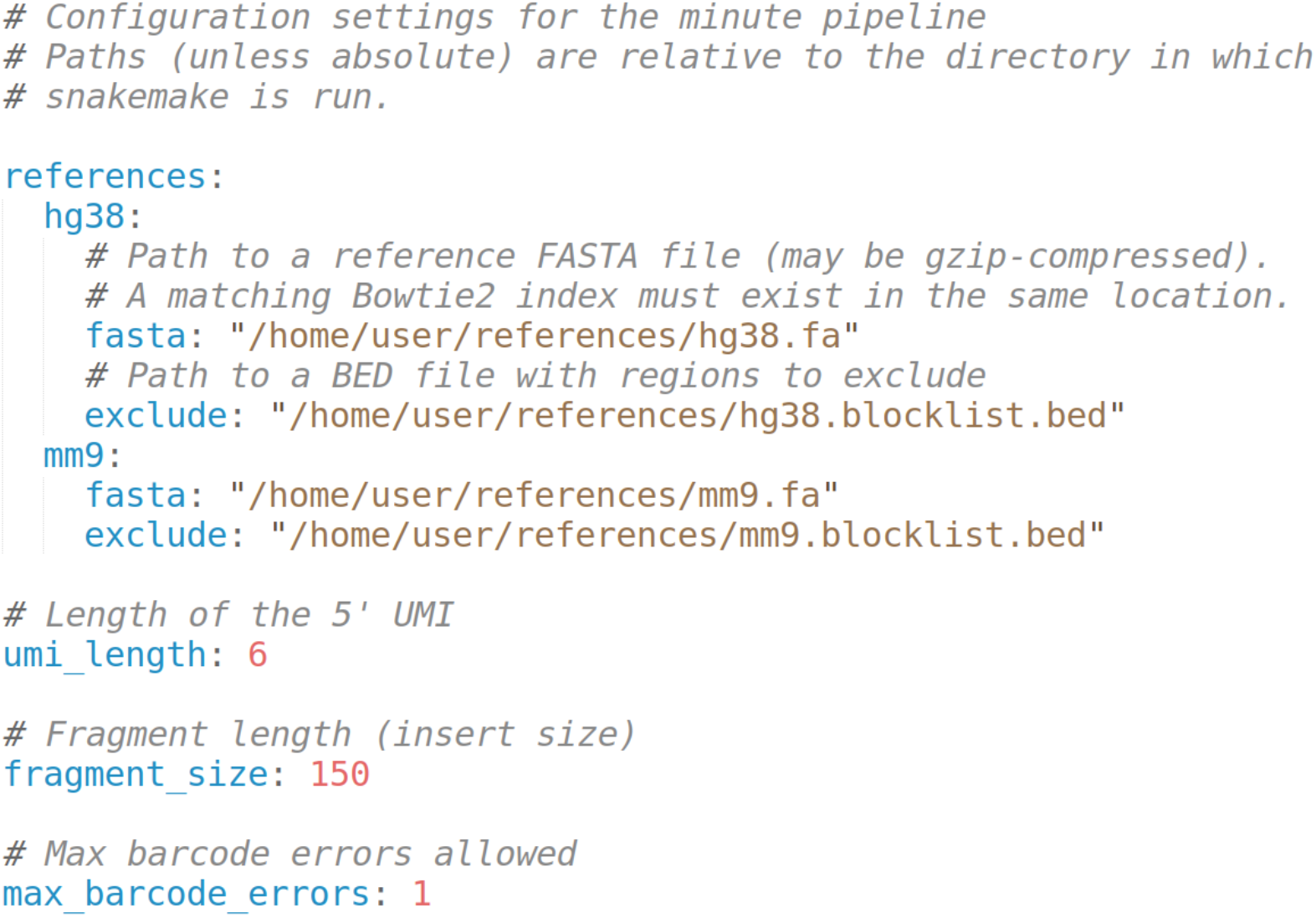
Example minute.yaml file. Reference genomes are configured specifying a path to a FASTA file and an optional exclude loci BED file. FASTA files must be indexed by bowtie2-index prior to running the pipeline. Any number of reference genomes can be included. The references specified in groups.tsv must match one of the references in minute.yaml. UMI length, fragment size and maximum barcode demultiplexing errors are also user-defined parameters.

**Figure 3.**
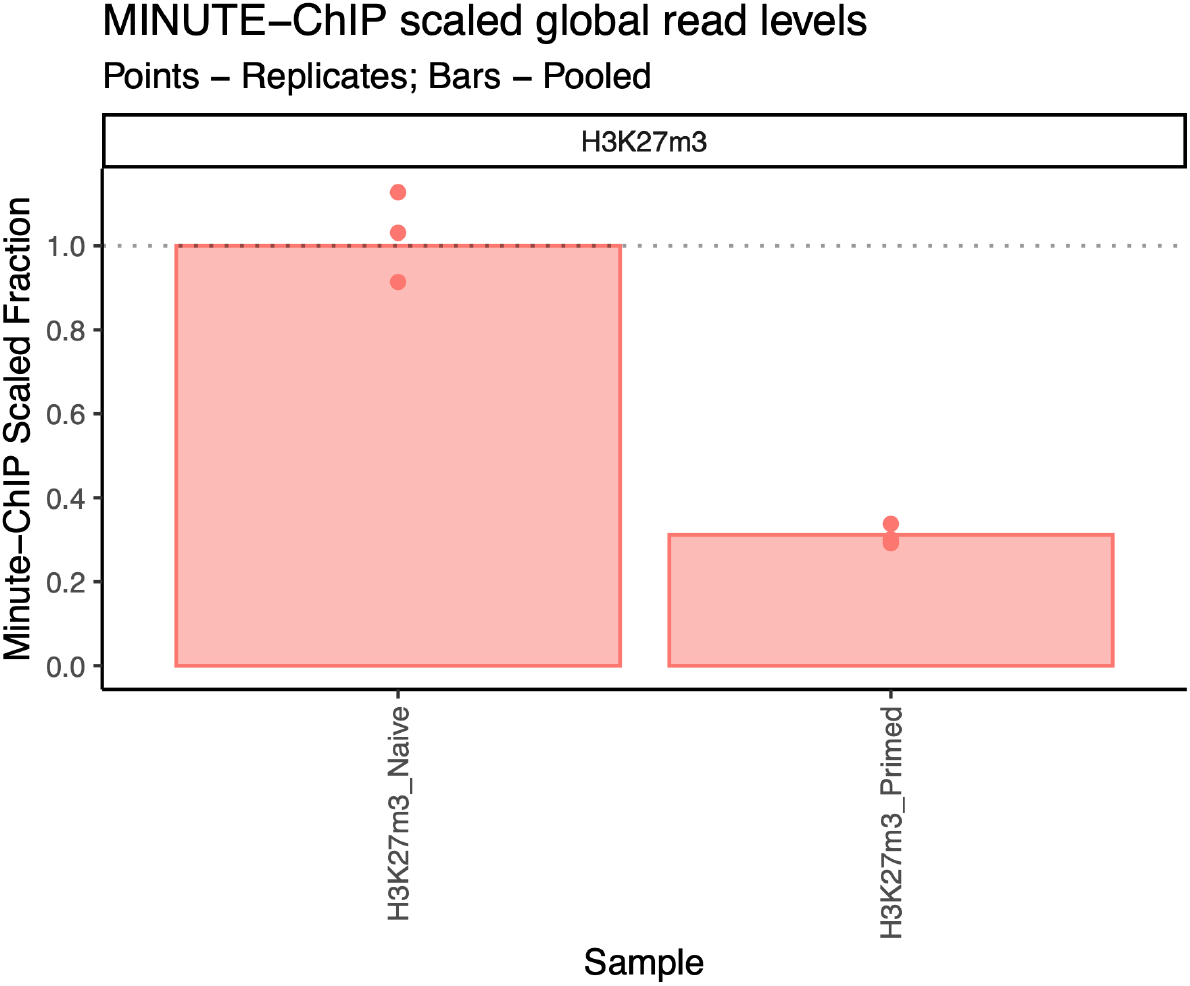
Global scaling of H3K27m3. Bars represent global ratio of pooled replicates and points represent each individual replicate. All values are scaled to H3K27m3 naïve hESCs pooled.

#### Output files

A minute pipeline run outputs i) a set of bigWig files, one unscaled file per condition (each line in the `libraries.tsv` file), and a scaled bigWig file per line in `groups.tsv`; ii) a matching deduplicated BAM alignment per library and iii) demultiplexed FASTQ files. Mapping statistics per library, scaling factors and a MultiQC (Ewels 2016) report summary is generated for the full run.

#### Run setup

Any minute experiment can be set up by the following steps:

1. Activate conda environment (see installation instructions): `conda activate minute`
2. Initialize a minute experiment using minute init: `minute init myexperiment --reads /path/to/fastq/`. This will automatically: Create a subfolder fastq in the `myexperiment` folder, create a fastq folder as a symbolic link to `/path/to/fastq` which should point to the input FASTQ files Read 1 and read 2 files must have names ending in `_R1.fastq.gz` and `_R2.fastq.gz`, respectively.
3. Edit minute.yaml file as needed.
4. Create `libraries.tsv` and `groups.tsv` files describing your libraries (see below) in the `myexperiment` folder;.
5. Run: `cd myexperiment; minute run` It is possible to run minute on High Performance Computing (HPC) and other environments. See minute and Snakemake documentation for more details.

## Example: **minute** run on a GEO dataset

Here, we illustrate the use of minute on a real dataset. To this end, we run minute on our H3K27m3 published dataset from (Kumar et al. 2021), available on GEO database. This dataset consists of H3K27m3 MINUTE-ChIP on naïve and primed human ES cells (hESCs). Note that GEO does not keep multiplexed FASTQ files, so the samples were submitted after demultiplexing. For this reason, minute can optionally skip the demultiplexing step when the barcode field in `libraries.tsv` is not provided (filled in with a. value). All configuration files described below can be found together as Supplementary file 1. In the following instructions, the path where Supplementary File 1 is decompressed to is called `myexperiment`.

### Download FASTQ files

minute has an auxiliary subcommand to download data from GEO that works using only the libraries.tsv file, provided that GEO accession numbers are set as the FASTQ file names (see Table 3).

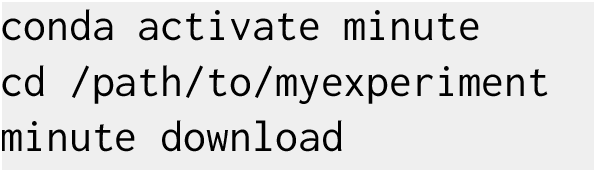

**Table 3.** libraries.tsv for the example run. H3K27m3 naïve and primed, three replicates each. Data from (Kumar et al. 2021) and available on GEO database for download under the corresponding accessing numbers. Barcodes are entered as “.”, since GEO data are already demultiplexed.

|  |  |  |  |
| --- | --- | --- | --- |
| H3K27m3_Naive | 1 | . | SRR15309374 |
| H3K27m3_Naive | 2 | . | SRR15309375 |
| H3K27m3_Naive | 3 | . | SRR15309376 |
| H3K27m3_Primed | 1 | . | SRR15309380 |
| H3K27m3_Primed | 2 | . | SRR15309381 |
| H3K27m3_Primed | 3 | . | SRR15309382 |
| Input_Naive | 1 | . | SRR17657859 |
| Input_Naive | 2 | . | SRR17657860 |
| Input_Naive | 3 | . | SRR17657861 |
| Input_Primed | 1 | . | SRR17657865 |
| Input_Primed | 2 | . | SRR17657866 |
| Input_Primed | 3 | . | SRR17657867 |

After the download run is complete, a `fastq/` directory with the corresponding FASTQ files must have been created. Alternatively, the user can manually download the FASTQ files from GEO. As long as the accession identifiers are kept as SRRXXXXXXXX_R1.fastq.gz, SRRXXXXXXXXX_R2.fastq.gz, the following steps work the same way.

### Specify experimental design

`libraries.tsv` file should contain four libraries with three replicates each: H3K27m3 on naïve and primed hESCs, plus their corresponding inputs (see Table 3). Additionally, `groups.tsv` contains the scaling and reference information (Table 4). Samples will be scaled relative to the combined H3K27m3 naïve samples (hence average of the three naïve replicates) as reference.

**Table 4.** groups.tsv for the example run. Each H3K27m3 library (first column) is matched to the corresponding input (third column). Pooled samples are added (second column) for robust normalization. Scaling group is named H3K27m3_Naive (4th column) and reads are mapped to hg38 reference (5th column).

|  |  |  |  |  |
| --- | --- | --- | --- | --- |
| H3K27m3_Naive | pooled | Input_Naive | H3K27m3_Naive | hg38 |
| H3K27m3_Naive | 1 | Input_Naive | H3K27m3_Naive | hg38 |
| H3K27m3_Naive | 2 | Input_Naive | H3K27m3_Naive | hg38 |
| H3K27m3_Naive | 3 | Input_Naive | H3K27m3_Naive | hg38 |
| H3K27m3_Primed | pooled | Input_Primed | H3K27m3_Naive | hg38 |
| H3K27m3_Primed | 1 | Input_Primed | H3K27m3_Naive | hg38 |
| H3K27m3_Primed | 2 | Input_Primed | H3K27m3_Naive | hg38 |
| H3K27m3_Primed | 3 | Input_Primed | H3K27m3_Naive | hg38 |

### Specify run configuration

`minute.yaml` (see Figure 2) contains paths to genome reference files and exclude BED annotations. Additionally, UMI length can be specified and a fragment length estimate is used to compute scaling factors.

Once all FASTQ files are downloaded and configuration setup is completed, the user needs to run the pipeline from the same `myexperiment/` directory where the `fastq/` directory is.

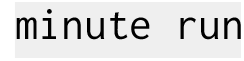

Once the run is complete, a final directory will contain three directories:

1. `bigwig.` Generated bigWig files, both scaled and unscaled (1x genome coverage) named after `libraries.tsv,` plus replicate number (or pooled) and the reference they were mapped to (e.g `H3K27m3_Naive_pooled.hg38.scaled.bw`, `Input_Primed_rep1.hg38.unscaled.bw`).
2. `bam.` Alignment BAM files for each library.
3. `demultiplexed.` Demultiplexed FASTQ files. In order to save disk space, in the specific case where the input FASTQ files are already demultiplexed, this directory contains symbolic links to the initial files.

Additionally, a reports directory is created with a MultiQC summary of all the statistics generated and minute scaling information. In this dataset, scaling shows that H3K27m3 global levels are about ~3 times higher in naïve than in primed cells (Figure 3).

BigWig files generated by minute can serve as a basis for further downstream analysis. Both scaled and unscaled files are provided for all libraries specified in the groups.tsv file, in order to provide comprehensive information about the influence of scaling on the global results. Visual inspection of these files on IGV already shows that high global levels of H3K27m3 in naïve hESCs can produce misleading results on unscaled data (Figure 4).

**Figure 4.**
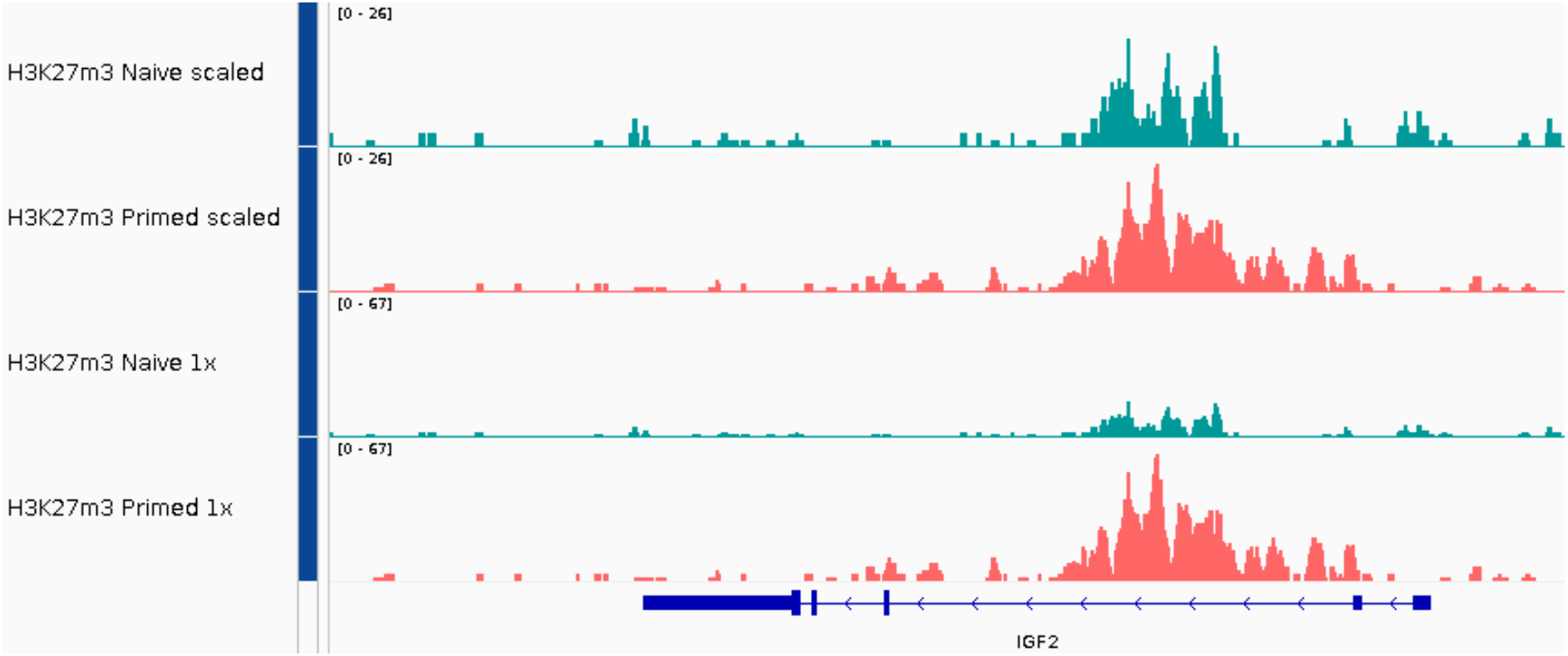
IGV view of IGF2, a gene that is affected by H3K27m3 scaling.

## Discussion

Minute is an open-source workflow that meets code quality standards required for robust processing of MINUTE-ChIP data. With very reduced user intervention, raw multiplexed FASTQ files from a MINUTE-ChIP experiment are transformed into a set of scaled, directly comparable bigWig files. Input genome tracks are also generated, along with a set of quality control metrics and relevant statistics reports. minute is implemented in Python and Snakemake, a well established workflow management system, and benefits from the advantages that such an engine provides, such as enhanced reproducibility, scalability, reentrancy and portability, among others. minute can be run on a variety of environments including high-performance computing, cloud and local environments. The resulting bigWig files can be further used for downstream analysis or visualization using genome browsing tools. As it has been shown in our previous studies (Kumar and Elsässer, 2019); Kumar et al. 2021), accurate global scaling of bigWig tracks can be crucial to biological discoveries that are overlooked with traditional non-quantitative ChIP-seq approaches.

## Supporting information

ZIP

## Data and source code availability

All FASTQ files used in the sample use case are available on Gene Expression Omnibus (GEO) database. Configuration files necessary for downloading them from GEO and running minute are also available as supplementary material.

Source code of minute is available on GitHub: https://github.com/NBISweden/minute

## Acknowledgements

We acknowledge the Swedish National Infrastructure for Computing (SNIC) at Uppmax server (projects SNIC 2020/15-9, SNIC 2020/6-3) for providing a high-performance computing environment to run minute software. MM is financially supported by the Knut and Alice Wallenberg Foundation as part of the National Bioinformatics Infrastructure Sweden at SciLifeLab. We thank members of the Elsässer lab and Manfred Grabherr for feedback on the pipeline development.

## Notes

### Competing Interest Statement

The authors have declared no competing interest.

