## Supplementary material for "minute: A MINUTE-ChIP data analysis workflow": ZIP: minute.yaml

```
# Sample minute.yaml config configuration settings for the minute pipeline
# Paths (unless absolute) are relative to the directory in which snakemake is
# run.

references:
  hg38: # Arbitrary name for this reference. This is used in output file names.
    # Path to a reference FASTA file (may be gzip-compressed).
    # A matching Bowtie2 index must exist in the same location.
    fasta: "/home/user/references/hg38.fa"
    # Path to a BED file with regions to exclude
    exclude: "/home/user/references/hg38.blocklist.bed"
  mm9:  
    fasta: "/home/user/references/mm9.fa"
    exclude: "/home/user/references/mm9.blocklist.bed"

# Length of the 5' UMI
umi_length: 6

# Fragment length (insert size)
fragment_size: 150

# Max barcode errors allowed
max_barcode_errors: 1
```
